# Does improved (Limu) maize variety adoption improves rural households’ food security: evidence from Dale Wabara district, Ethiopia

**DOI:** 10.1101/2023.06.06.543866

**Authors:** Bikila Tesfa, Adugna Eneyew, Fikadu Mitiku

## Abstract

Adopting improved agricultural technology is important instruments to ensure household food security. There is, however, limited empirical evidence on impacts of adoption of improved maize varieties grown by smallholder farmers. Consequently, the study examined the impact of Limu maize variety adoption on rural households’ food security in Dale Wabara District, Ethiopia. Quantitative and qualitative data were collected from both primary and secondary sources. Three-stage sampling technique was employed to select 319 households for primary data collection. Binary logit model regression model and propensity score matching model were used to investigate determinants of Limu maize variety adoption and its impact on rural households’ food security, respectively. Furthermore, daily calorie intake per adult equivalent was used to measure food security status of household in study area. According to the results of the binary logit regression model households head education, family size, total land size, frequency of extension visits, livestock ownership and availability of Limu maize seed increases the likelihood of household’s adoption of Limu maize variety. The result of propensity score matching model shows that adoption of improved Limu maize variety significantly improve household food security (daily calorie intake). Empirical evidence shows that adopter households are better-off than non-adopter households in terms of their daily calorie intake. Thus, it can be concluded that Limu maize variety adoptions significantly contribute to the economic and social development of smallholder farmers by increasing food security status in study areas. It is therefore recommended that wider distribution of Limu maize variety has to be prioritized to improve food security in study areas.

## 1. Introduction

Maize (*Zeameys*) is the most extensively cultivated crop in Ethiopia, which is produced primarily using rainwater under a variety of agroecological and socioeconomic situations (1). The third-largest producer of maize in Africa is Ethiopia. About 2.13 million hectares of land were covered with maize during the 2017–2018 growing season, yielding 83.96 million quintals (2).

The main food source and a crucial component of Ethiopian households’ food security is maize (1). It is food source for more than 80% of Ethiopians (3). The small-scale farmers themselves use three-fourths of the maize produced at the home level (2). No other grain crop that has been grown attains this degree of retention for domestic use. As a result, the amount of maize grain produced by smallholder farmers in a given year directly affects how people perceive their level of food security (4). As a result, in Ethiopia, maize is a crucial crop for food security.

Despite being a crucial crop for food security, Ethiopia’s average production of maize in 2018 was only 3.6 tons per hectare, which is lower than the global average of 5.6 tons per hectare (2). Therefore, to increase maize yield, the Ethiopian government has supported technology-driven programs. One of the agricultural policies supported to increase maize output and productivity as well as to better the food security situation of the maize producer farmers is the improvement of maize technology through the facilitation of research systems. The government also took steps to generate maize types that provide a high yield and are resistant to disease. As a result, more than 40 maize varieties, including hybrids and open-pollinated varieties (OPVs), have been generated for the country’s various agroecology (4).

Maize is one of the main cereal crops that is prioritized as a way to improve the life of the rural population, and the Dale Wabara district is one of the prospective maize producing districts in the Kellem Wollega zone. However, the low maize yield issue is a problem for farmers in the research area. Before 20 years ago, farmers began utilizing better maize types to lessen this issue. The productivity and production of maize, which affect rural peoples’ livelihoods, are still quite low. For example, during the 2017–18 growing season, maize covered 3,236 ha of land, of which 2,023 ha and 1213 ha were planted with traditional varieties and improved varieties, respectively, yielding an average of 20 and 36 quintals per hectare (5). The yield gap between local and improved maize variety was 16 quintals per ha which is substantial for achieving food security.

In terms of land coverage, 62.5% of the land allocated for maize production is covered by the local maize varieties and 37.5% by the improved maize varieties. This shows that the low productivity of maize is linked to the lower utilization of improved maize variety among maize producing farmers.

Some studies were conducted to investigate the impact of adopting improved maize varieties (6) (7) (8). There is, however, scant research on how improved maize variety adoption in Ethiopia may affect food security. Additionally, the majority of them were carried out at a macro level, which could make the results difficult to generalize to a particular location. Furthermore, to the researchers’ knowledge, no research has been done on how the adoption of the Limu maize variety affects the food security of rural households in the study area. Hence, it was not well known to what extent Limu maize variety adopter households are better off than non-adopter households in terms of their food consumption. Given the aforementioned rationale, the concept of Limu maize variety adoptions on food security in the study area needs to be operationalized. so that a common understanding could be reached. Therefore, this study generates evidence to fill the gaps of past studies by investigating the impact of Limu maize variety adoption on rural households’ food security in the Dale Wabara district, Ethiopia. Generally, this study has two contributions. First, it generates evidence on the effect of the Limu maize variety adoption on households’ food security. Second, it identifies factors influencing the adoption of improved Limu maize varieties. So that policymakers and local agricultural development planners use the information to improve farmers’ adoption of improved maize varieties. In addition, the finding from this study could also help agricultural policymakers elsewhere in developing countries with similar contexts particularly Sub-Saharan Africa.

## 2. Research Methodology

### 2.1. Study area

The study was conducted at Dale Wabara District of Kellem Wollega Zone, Oromia National Regional State of western Ethiopia. The district has 22 rural and 2 town *Kebeles* (the lower administrative level in Ethiopia) administrations with a total of 24 *Kebele* administrations. Dale Wabara district is located at 549Km distance from Addis Ababa, along the direction of western Ethiopia and lies within the Abay Valley drainage system (9).

### 2.2. Data collection method

Primary data were collected through various data collection instruments such as household survey, Focus Group Discussion and Key Informant interview. Focus group discussion and a key informant interview were also used to get additional supporting qualitative evidence on current situation of household food security and challenges that farmers have been faced in adoption of Limu maize variety. Data were collected from both primary and secondary sources on a wide variety of variables. The primary data was collected through individual interviews of selected respondents and survey was administered using structured questionnaires by trained enumerators who collected data from households through personal interviews. The survey collected information on several factors including household demographic characteristics (family sizes, age, sex and education level, etc.), socioeconomic factors and institutional factors were included in the data. To complement the primary data, secondary data were obtained from different unpublished and archival sources such as articles, official reports of relevant stakeholders, CSA report data and personal coummunications.

### 2.3. Sampling procedure

The study was done based on cross-sectional data. Three-stage sampling technique were applied to select sample households. In the first stage, four *Kebele* administrations (Kara Jeno, Meki Dimbar, Sago Adami and Dogano Bile) were selected randomly. In the second stage, maize producing farmers were stratified in to adopter and non-adopter farmers. Finally, farm household were selected using systematic random sampling technique by taking in to account proportional to the size of the population in each kebele administrations. The sample size of the households for this study was determined using the simplified formula provided by Yamane (10) to determine the required sample size at 95% confidence level.

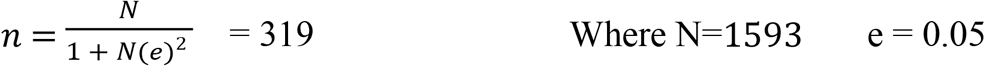

### 2.4. Data analysis

For data analysis, descriptive statistics such as mean, percentage, and standard deviations) were applied. While chi square (discrete variables) and t test (continuous variables) was applied to test the statistical significance association and difference among adopters and non-adopters, respectively. On behalf of econometrics model, binary logistic regression and propensity score matching model were employed. Binary logistic regression model was used to identify major determinants of adoption of Limu maize variety. Propensity score matching (PSM) was applied to investigate the impact of Limu maize variety adoption on rural households’ food security. Moreover, qualitative data collected from focus group discussion and key informant interview were analyzed by narrative explanation to complement quantitative data. Finally, the parameters were estimated by the maximum likelihood technique with the help of (SPSS ver. 20, STATA version 14 and Ms excel).

### 2.5. Empirical model specification

Binary logistic logistic regression model is a proper model when the dependent variable is a dummy one consisting of two, 0 and 1. Thus, logistic regression model that was employed is a binary logistic regression model, where dependent variable is Y and dependent one is X. Therefore, to identify the major determinants of adoption of Limu maize variety, binary logistic regression was used. In order to explain the model, the following logistic distribution function was used (11).

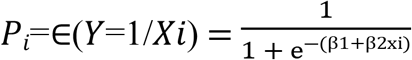

In the logistic distribution equation, *P*_*i*_ is the independent variable; *X*_i_ is the data that is the possibility of preference by an individual (option of having 1 and 0 values)

To estimate PSM either of the binary logistic regression and probit model can be used. However, binary logistic regression has been chosen. In binary logistic regression model, the slope coefficients of a variable give the change in the log of the odds associated with a unit change in the variable, holding all other variable constant. The rate of change in the probability of an event happening is given by βjPi(1-Pi), where βj is partial regression coefficient of the j^th^ regressor. But in evaluating Pi, all the variables included in the analysis are involved (11).

### 2.6. Impact assessment approach

According to Khandker (12) impact evaluation is the act of studying whether the changes in food security are indeed due to the adoption of improved maize variety. There are several approaches, which can be used to evaluate impacts. These include randomized selection methods, PSM, regression discontinuity design, the difference in difference, and instrumental variable estimation methods (13). There are numerous robust methods in the study of impact of a particular technology such as improved seeds. For instance, the randomized experimental method can be used to assess the impact of program, when a participant is randomly selected for it. Moreover, the difference method can be also be used when baseline and time-series information on both participants and non-participants is available (14). However, this study is cross sectional study where there are no randomized data or base line data. Therefore, in this study PSM method was used because PSM is the appropriate method when such kind of problem arises (13).

Propensity score matching constructs a statistical comparison group based on a model of the probability of adopting Limu maize variety, using observed characteristics. Then adopter households were matched to the non-adopter households based on the probability of the adopting the Limu maize variety. Since matching adopters to non-adopters on each covariate were practically difficult, propensity score was predicted based on observed characteristics of adopters and non-adopters. Then, the average treatment effect of the Limu maize variety adoption was calculated as the mean difference in outcomes between these two groups (15).

To estimate the average treatment effect on treated (ATT) by using the PSM method the steps such as estimation of the propensity scores, choosing a matching algorithm, checking on common support region, testing the matching balance and sensitivity analysis were followed.

### 2.7. Variables Selection, Definition and its Measurement

#### Adoption decision

The dependent variable has dichotomous in nature representing the farmers’ adoption decision of Limu maize variety; taking values, 1 for household who adopt Limu maize variety and 0 otherwise. In this paper a farmer household is categorized as an adopter if she/he used Limu maize variety in the last three years from 2019 to 2021 productions seasons and non-adopters are a household who have not used these seed in the last three years in the same period.

#### Outcome variable

daily calorie intake per adult equivalent is the outcome variable of the study. To measure the food security status of households in the study area, information concerning the type and amount of foods items consumed by each household in the seven days preceding the survey were collected from both adopter and non-adopter households. Then, the calorie content of food items consumed by sample households was calculated using the calorie conversion factor per adult equivalent. Finally, the amount of total calories consumption of each sample households was computed and divided by seven days and to Adult Equivalent (AE). This method measures food consumption directly and not only food availability. In addition, it addresses both dietary quality and caloric intakes at the household level (16). This is the reason why daily calorie intake per adult equivalent has been selected over the other.

## 3. Results and Discussion

### 3.1. Limu maize variety adoption status

The results of the study shows that 48.9% of respondent farmers in the study area had adopted the Limu maize variety. Comparing with previous varietal adoption studies in Ethiopia, this result shows that, increased generation and dissemination of improved maize technologies by government and non-government organizations as well as utilization of improved maize seed by maize growers in the past years. For instance, according to a survey conducted in 2013 by the Ethiopian Institute of Agricultural Research in collaboration with the International Maize and Wheat Improvement Center about 31% of the sampled farmers planted improved maize varieties (17).

Further, a study by Jaleta (7) revealed that the adoption status of improved varieties by households was 27% in 2016 in Ethiopia. The result of the current study was more than the previous study due to the study area is well known in maize production potential even from Ethiopia. In addition, maize is potential crop next to coffee and hence, prioritized as a means to improve the livelihood of rural community in the study area. This may the reason why adoption status of Limu maize variety adoption was high in the study area.

### 3.2. Food security status of households

Descriptive result of the outcome variable food security status of sample household measured in daily kilo calorie intake per adult equivalent. The average adult equivalent daily calorie intake for adopter and non-adopter households is 2555.87 and 2082.67, respectively. This demonstrates unequivocally that adoptive families consumed 473.2 Kilocalorie (kcal) more on average than non-adopter households did. In terms of kcal intake per adult equivalent, adopter and non-adopter households differ significantly from one another. Based on the 2200 kcal/day/adult equivalent cut point, about 52.35 percent of respondent homes in the research area had food insecurity, compared to approximately 47.65 percent of respondent households who had food security.

### 3.3. Determinants of improved (Limu) maize variety adoption from the estimation of binary logistic regression

The estimated binary logistic regression model indicated that six (6) of the twelve (13) explanatory variables significantly influenced Limu maize variety adoption (Table 2). Those are:

**Table 1:** Summary of Variables selection, definition and its measurement

| No | Variables | Type of Variable | Measurement & definition | Hypothesis |
| --- | --- | --- | --- | --- |
| I | Limu maize variety adoption | Dummy | 1 if adopters otherwise 0 |  |
| II | Outcome variable | Continuous | Households' daily calorie intake per adult equivalent |  |
| 1 | Gender | Dummy | 1 if male, otherwise 0 | + |
| 2 | Age of HH | Continuous | Year | - |
| 3 | Education of HH | Continuous | Year of schooling | + |
| 4 | Family size | Continuous | Adult equivalent | + |
| 5 | off/non-farm income | Dummy | 1 if yes or 0 otherwise | + |
| 6 | Cooperative membership | Dummy | 1 if yes or otherwise 0 | + |
| 7 | Access to credit | Dummy | 1 if yes or 0 otherwise | + |
| 8 | Frequency of Extension contact | Continuous | Number per year | + |
| 9 | Distance to the market | Continuous | Kilometer | - |
| 10 | Total livestock ownership | Continuous | Tropical Livestock Unit | + |
| 11 | Total land size | Continuous | Hectare | + |
| 12 | Lag Price of Maize Grain | Continuous | Ethiopian Birr | + |
| 13 | Availability of improved maize seed | Dummy | 1 if available or 0 otherwise | + |

**Table 2:**
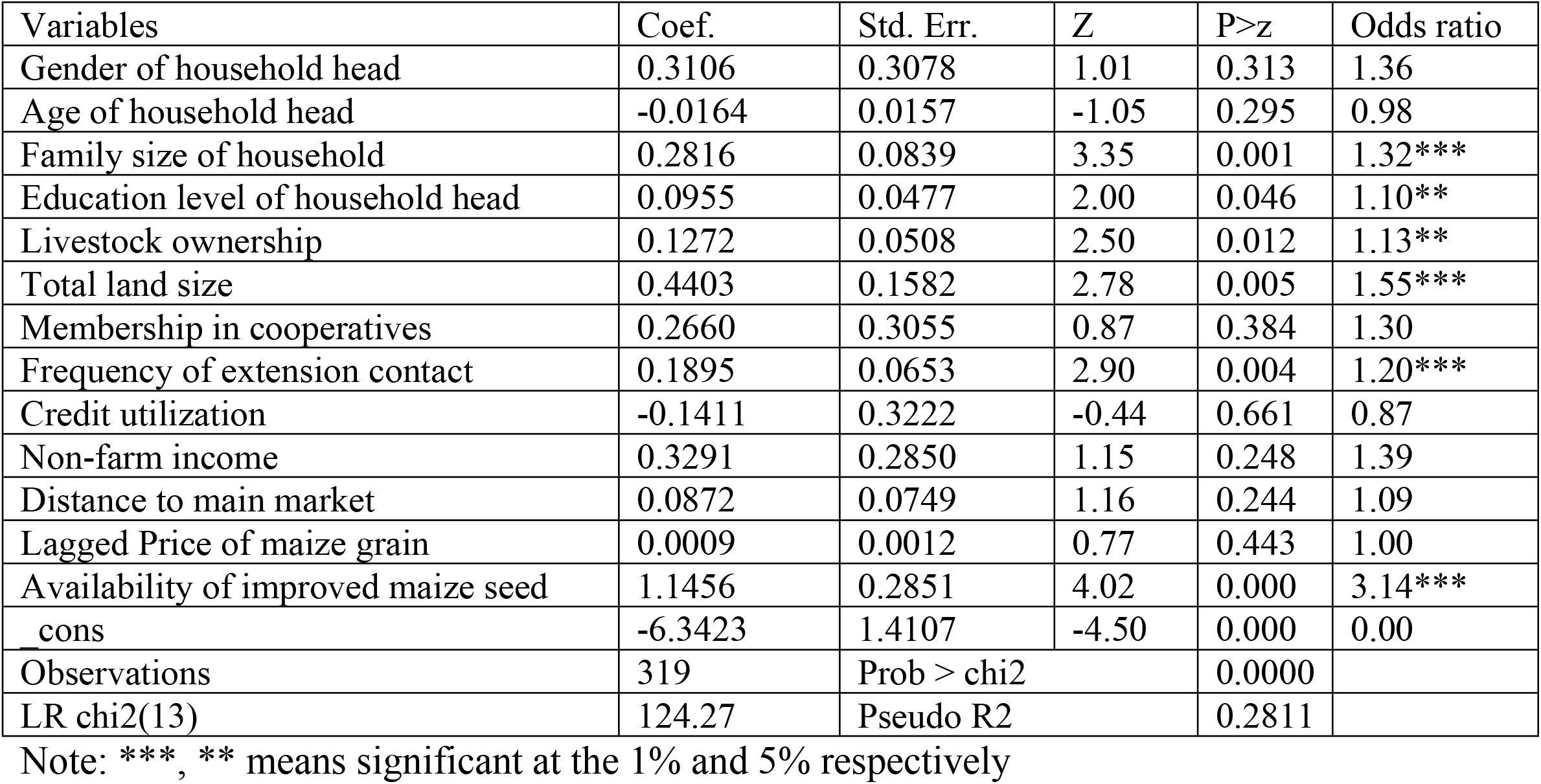
Binary logistic regression model results of Limu maize variety adoption

| Variables | Coef. | Std. Err. | Z | P>z | Odds ratio |
| --- | --- | --- | --- | --- | --- |
| Gender of household head | 0.3106 | 0.3078 | 1.01 | 0.313 | 1.36 |
| Age of household head | -0.0164 | 0.0157 | -1.05 | 0.295 | 0.98 |
| Family size of household | 0.2816 | 0.0839 | 3.35 | 0.001 | 1.32*** |
| Education level of household head | 0.0955 | 0.0477 | 2.00 | 0.046 | 1.10** |
| Livestock ownership | 0.1272 | 0.0508 | 2.50 | 0.012 | 1.13** |
| Total land size | 0.4403 | 0.1582 | 2.78 | 0.005 | 1.55*** |
| Membership in cooperatives | 0.2660 | 0.3055 | 0.87 | 0.384 | 1.30 |
| Frequency of extension contact | 0.1895 | 0.0653 | 2.90 | 0.004 | 1.20*** |
| Credit utilization | -0.1411 | 0.3222 | -0.44 | 0.661 | 0.87 |
| Non-farm income | 0.3291 | 0.2850 | 1.15 | 0.248 | 1.39 |
| Distance to main market | 0.0872 | 0.0749 | 1.16 | 0.244 | 1.09 |
| Lagged Price of maize grain | 0.0009 | 0.0012 | 0.77 | 0.443 | 1.00 |
| Availability of improved maize seed | 1.1456 | 0.2851 | 4.02 | 0.000 | 3.14*** |
| cons | -6.3423 | 1.4107 | -4.50 | 0.000 | 0.00 |
| Observations | 319 | Prob > chi2 |  | 0.0000 |  |
| LR chi2(13) | 124.27 | Pseudo R2 |  | 0.2811 |  |

Note: \*\*\*, \*\* means significant at the 1% and 5% respectively

#### Family size

this variable significantly affects Limu maize variety adoption positively as expected at 1% significant level (table 2). This shows that having a large family gives farmers additional labor for tasks like planting, drilling, and weeding as well as more incentive to grow more maize grain. A further factor increasing the significance of family size as a source of labor in the adoption of the Limu maize variety is the fact that agriculture in the study area is a labor incentive. This result is also in agreement with previous empirical findings such as (6, 18).

#### Education level of household head

as expected, education level of the household head positively and significantly influences the adoption of Limu maize variety at 5% significance level (table 2). This may be because education increases awareness, encourages farmers to embrace the Limu maize variety, and teaches them to comprehend and analyze various pieces of information about emerging technologies. This result is in line with the finding of (7, 19, 20).

#### Livestock ownership

as hypothesized, this variable significantly and positively affects adoption of Limu variety at 5% significance level (table 2). This may as a result of the fact that livestock are a significant source of food, draft power, and additional cash that helps farmers to purchase Limu maize seed. This is in line with the study by (6, 21) which confirms the positive relationship between livestock and adoption.

#### Total land size

The result is as expected, total land size has a positive and significant relationship at the 1% significant level (table 2). This supports the hypothesis that having more hectare of land is associated with higher adoption rates of the Limu maize variety. Due to the dependence of rural residents’ livelihood on land, it is one of the most crucial inputs for agricultural production. This result is consistent with findings such as (7, 20, 22, 23).

#### Availability of improved maize seed

as proposed, availability of improved maize seed positively and significantly influences the probability of Limu maize variety adoption at 1% significant level (table 2). This implies that availability of technology is bottleneck factor for adoption of new technology. Moreover, the farmers during the FGD session confirm that improved maize seed and fertilizer in study area is not available at requested time and amount. In addition, improved maize seed and fertilizer is arrived after time of plantation. A study by Spielman (24) also confirms that the timely deliverance of improved seed affects the utilization of improved seed due to incompatibility of cropping season and time of deliverance. This finding is also in line with (25, 26) which confirmed that unavailability of the required seeds at the right place and time are key factors that affecting improved seed adoption.

#### Frequency of extension contacts

the binary logistic regression model output shows that the households’ frequency of extension contact per year had positively and significantly influenced the likelihood of adoption of Limu maize variety at a 1% significance level (table 2). Since extension contacts are the primary source of agricultural knowledge in Ethiopia, they are a crucial catalyst for technology adoption. The results of the key informant interview showed that farmers who had greater interaction with extension agents were encouraged to attend farmers’ training; this enhanced their knowledge and skills on agricultural methods, resulting in an improvement in the use of improved maize varieties. This demonstrates that getting instruction from development agents and their counsel, as well as the perceived value of their guidance, are key elements in explaining the likelihood of Limu maize variety adoption. This finding of this research is also in lined with the research result reported by (18, 19, 27). On contrary, (28) report a negative and significant relationship between access to extension measured by the number of contacts a farmer had with extension agents and the likelihood of using improved maize varieties.

### 3.4. Impact of Limu maize variety adoption on rural households’ food security

#### 3.4.1. Estimation of Propensity Score

The coefficients of the binary logistic regression model have been used to generate propensity scores that can be further used for matching purposes. Based on the estimated propensity scores, 156 adopter households have been matched to 163 non-adopter households (control groups) that most resemble them. Twelve matching variables have been used in the model as explanatory variables. In doing so, the dependent variable was a binary variable taking a value of 1 for adopter household or otherwise. Results presented in table 2 shows the estimated model appears to perform well for the intended matching exercise. The rationale for PSM helps to compare adopters Limu maize variety with non-adopter households. Additionally, pseudo-R2 has been computed. The pseudo-R2 value represents how well the regressors explain the likelihood of involvement. A reliable estimate of the model requires a low value for the pseudo (29). Hence, the model is considered good since the estimated actual pseudo-R^2^ is 0.2690.

#### 3.4.2. Choosing matching algorithm

The selection of a matching algorithm is the following step in the ATT calculation. Different matching estimators were tested in the common support region to match adopter households of the Limu maize variety with non-adopter households. Different factors, such as an equal means test (balance test), a low pseudo-R2 for the overall balancing test, and a higher number of matched sample sizes, are used to choose the best matching estimator (30). Balancing test refers to the explanatory variables with no statistically significant mean differences between the matched groups of treated control households. Based on equal means testing (balance test), a low pseudo-R2 for the overall balancing test, and a larger number of matched sample sizes, the results show that kernel matching with a band width of (0.25) is the best estimator for the data at hand (Table 3). Consequently, the estimation outcomes and discussion below are the immediate results of the kernel matching with band width of (0.25).

**Table 3:** Performance criteria of matching algorithms

| Matching estimators | Balancing test* | Pseudo-R<sup>2</sup> after matching | Matched sample size |
| --- | --- | --- | --- |
| Nearest Neighbor (NN) |  |  |  |
| Neighbor (1) | 8 | 0.056 | 312 |
| Neighbor (2) | 10 | 0.050 | 312 |
| Neighbor (3) | 13 | 0.029 | 312 |
| Neighbor (4) | 12 | 0.022 | 312 |
| Neighbor (5) | 13 | 0.020 | 312 |
| Caliper matching (CM) |  |  |  |
| 0.01 | 13 | 0.025 | 288 |
| 0.05 | 8 | 0.056 | 312 |
| 0.1 | 8 | 0.056 | 312 |
| 0.5 | 8 | 0.056 | 314 |
| Kernel Matching (KM) |  |  |  |
| With band width of (0.08) | 13 | 0.020 | 312 |
| With band width of (0.1) | 13 | 0.018 | 312 |
| With band width of (0.25) | 13 | 0.014 | 312 |
| With band width of (0.5) | 7 | 0.059 | 312 |
| Radius Matching |  |  |  |
| With band width of (0.01) | 6 | 0.273 | 312 |
| With band width of (0.1) | 6 | 0.273 | 312 |
| With band width of (0.25) | 6 | 0.273 | 312 |
| With band width of (0.5) | 6 | 0.273 | 312 |

#### 3.4.3. Identifying common support condition

The region with the lowest and highest propensity scores for the treatment and control groups of families, respectively, is known as the common support region. A common support condition is imposed to ensure that any combinations of traits seen in the treatment group can also be seen in the control group (31). The analysis should only include the subset of the comparison group that is comparable to the treatment group; any findings outside of this should be disregarded (32). According to table 4 below, the propensity score for families that have adopted the Limu maize variety ranges from 0.055 to 0.983 with a mean score of 0.665, while it ranges from 0.012 to 0.963 for households that have not adopted the variety, with a mean score of 0.320. so between 0.05 and 0.96 is where there is a common support. Therefore, for matching purposes, households with propensity scores between 0.055 and 0.963 are not taken into account. This is due to the fact that when there is no overlap between the treatment and non-treatment, no matches can be established to estimate the average treatment effects on the ATT parameter (31). For the sake of this restriction, 7 households all from adopter were discarded. This shows that the study does not have to drop many adopter households from the sample in computing impact estimator.

**Table 4:** Distribution of Estimated propensity score of households

| Group | Observation | Mean | SD | Min | Max |
| --- | --- | --- | --- | --- | --- |
| All households | 319 | 0.489 | 0.292 | 0.0118 | 0.983 |
| Adopter households | 156 | 0.665 | 0.222 | 0.055 | 0.983 |
| Non-adopter Households | 163 | 0.320 | 0.248 | 0.012 | 0.963 |

As shown in table 5, the total treated observations 7 households (2.19%) are off-support, while 314 households (97.81%) are on support and all the control households are included in the common support region.

**Table 5:**
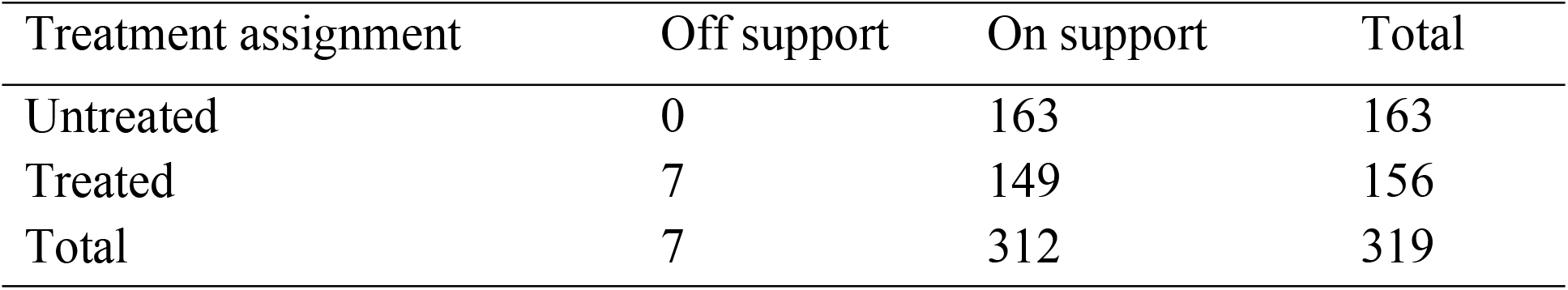
Common Support region

| Treatment assignment | Off support | On support | Total |
| --- | --- | --- | --- |
| Untreated | 0 | 163 | 163 |
| Treated | 7 | 149 | 156 |
| Total | 7 | 312 | 319 |

Testing to see if the propensity score effectively balances characteristics between the treatment and comparison group units is a crucial step in evaluating the quality of matching. The calculated propensity scores for adopter and non-adopter households are shown in the histogram in Figure 1 below. The common support criterion is met, as shown by the large overlap in the distribution of the propensity scores of both the adopter and non-adopter groups, according to a visual assessment of the density distributions of the estimated propensity scores for the two groups. The graph’s upper half pertains to adopters, while the bottom half depicts the propensity scores distribution for non-adopters. On the y-axis are the score densities.

**Figure 1:**
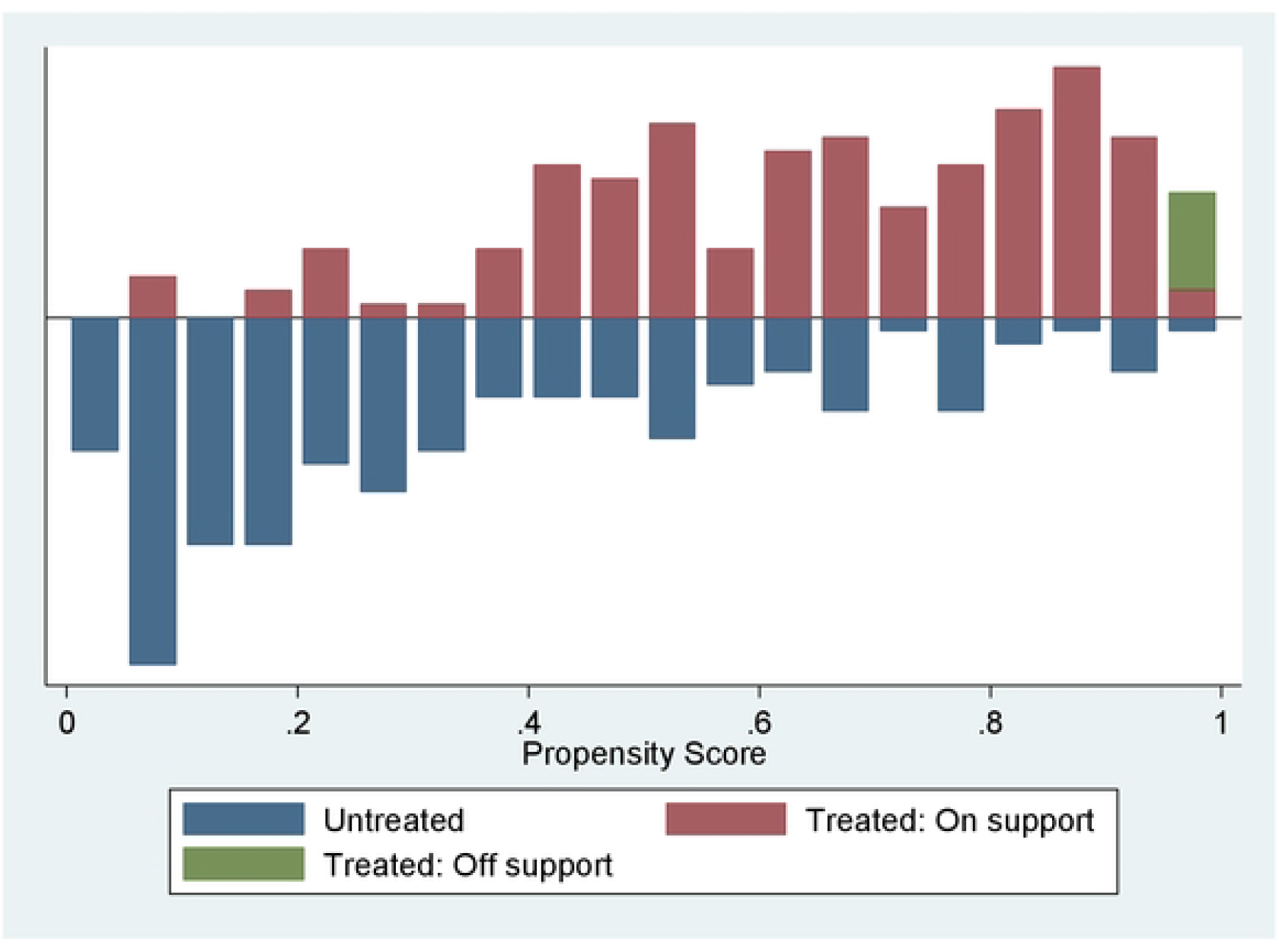
Graph of common support region

#### 3.4.4. Testing the balance of propensity score and covariates

The next stage, after selecting the optimal matching algorithm and common support condition, is to apply the chosen matching algorithm and check the propensity score and covariate balance using various approaches. Different test methodologies, such as the reductions in mean standardized bias between matched and mismatched households, equality of means using t-test and chi-square test for joint significance for the variables utilized, are taken into account when determining the balancing capacities of the estimations.

Following Rosenbaum (15), the standardized difference has been calculated, that is, the size difference in means of the conditioning variables (between adopters and non-adopters). In the present matching models, the standardized difference in Z before matching is in the range of 5.5% and 78.6% in absolute value. Rosenbaum and Rubin (15) recommend that a standardized difference after matching of 20% or more should be viewed as large. Accordingly, the remaining standardized difference of Z after matching for all covariates lies between 0.4 and 19% which is below the critical level of 20% (table 6). in all cases, it is evident that sample differences in the unmatched data significantly exceed those in the samples of matched cases. The process of matching thus creates a high degree of covariate balance between the treatment and control samples that are ready to use in the estimation procedure.

**Table 6:** Propensity score and covariate balance

| Variable | Unmatched | Mean |  | %bias | %reduct<br>bias | t-test |  |
| --- | --- | --- | --- | --- | --- | --- | --- |
|  | Matched | Treated | Control |  |  | t | p>t |
| Pscore | U | 0.66536 | 0.32027 | 146.5 |  | 13.06 | 0.000 |
|  | M | 0.65083 | 0.61312 | 16.0 | 89.1 | 1.47 | 0.141 |
| Gender of the household | U | 0.72436 | 0.69939 | 5.5 |  | 0.49 | 0.624 |
|  | M | 0.71141 | 0.69853 | 2.8 | 48.4 | 0.24 | 0.808 |
| Age of the household | U | 41.468 | 46.503 | -40.7 |  | -3.63 | 0.000 |
|  | M | 41.591 | 41.998 | -3.3 | 91.9 | -0.30 | 0.761 |
| Family size | U | 4.6813 | 3.6592 | 61.1 |  | 5.46 | 0.000 |
|  | M | 4.6036 | 4.2859 | 19.0 | 68.9 | 1.64 | 0.102 |
| Education level | U | 7.2244 | 4.3681 | 70.0 |  | 6.25 | 0.000 |
|  | M | 6.9396 | 6.6133 | 8.0 | 88.6 | 0.69 | 0.489 |
| Livestock ownership | U | 10.143 | 7.8803 | 70.2 |  | 6.26 | 0.000 |
|  | M | 9.9338 | 9.7064 | 7.1 | 90.0 | 0.60 | 0.546 |
| Total land size | U | 2.8013 | 2.0383 | 78.6 |  | 7.00 | 0.000 |
|  | M | 2.7852 | 2.7809 | 0.4 | 99.4 | 0.04 | 0.971 |
| Being member of cooperative | U | 0.72436 | 0.62577 | 21.1 |  | 1.88 | 0.061 |
|  | M | 0.71812 | 0.71921 | -0.2 | 98.9 | -0.02 | 0.983 |
| Frequency of extension contact | U | 3.9936 | 2.9448 | 48.8 |  | 4.36 | 0.000 |
|  | M | 3.8859 | 3.7726 | 5.3 | 89.2 | 0.44 | 0.659 |
| Access to credit | U | 0.73718 | 0.70552 | 7.0 |  | 0.63 | 0.530 |
|  | M | 0.73826 | 0.71729 | 4.7 | 33.8 | 0.41 | 0.686 |
| Non-farm income | U | 0.55128 | 0.51534 | 7.2 |  | 0.64 | 0.522 |
|  | M | 0.53691 | 0.52553 | 2.3 | 68.3 | 0.20 | 0.845 |
| Distance to main market | U | 7.5032 | 7.1871 | 16.8 |  | 1.50 | 0.135 |
|  | M | 7.4933 | 7.5622 | -3.7 | 78.2 | -0.33 | 0.741 |
| Lagged Price of maize grain | U | 723.72 | 706.44 | 14.6 |  | 1.30 | 0.193 |
|  | M | 725.84 | 723.61 | 1.9 | 87.1 | 0.16 | 0.870 |
| Availability of improved maize seed | U | 0.71795 | 0.41104 | 64.9 |  | 5.79 | 0.000 |
|  | M | 0.7047 | 0.69039 | 3.0 | 95.3 | 0.27 | 0.789 |

Kernel matching has been taken into account in this investigation. The outcome shows that some variables had statistically significant differences before matching. However, after matching, the factors are typically balanced and there are no discernible changes. The matching has been

deemed a valid match since the mean differences for all covariates between the treated and control groups after matching have not been determined to be statistically significant (or if the p-value calculated after matching is >0.05). As a result, valid matching has been provided for all covariates using kernel matching.

Table 7 shows that the standardized mean difference for all covariates utilized in the propensity score, which was approximately 46.6% before matching, is now only about 5.5%. The joint significance of the covariates was always rejected after matching, although it was never rejected before matching, according to the p-value of the likelihood ratio tests. Through matching, the bias was significantly decreased, falling in the 33 to 22% range. Additionally, the pseudo R2 was dramatically reduced from 28% prior to matching to roughly 1.4% after matching. This low pseudo R2, low standardized bias, high overall bias reduction, and the likelihood ratio test’s inconsequential p-values after matching all point to the propensity specification’s success in balancing the distribution of covariates.

**Table 7:**
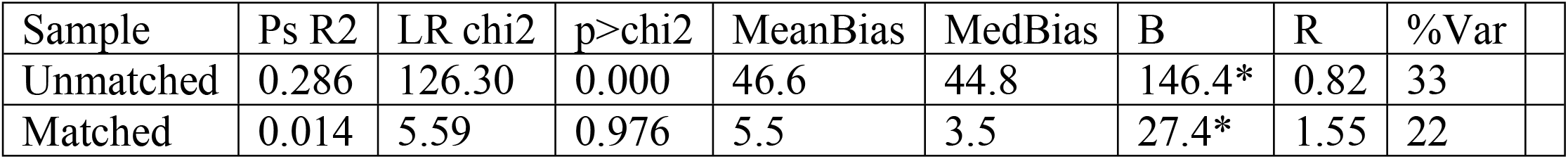
Propensity score matching: quality test

| Sample | Ps R2 | LR chi2 | p>chi2 | MeanBias | MedBias | B | R | %Var |
| --- | --- | --- | --- | --- | --- | --- | --- | --- |
| Unmatched | 0.286 | 126.30 | 0.000 | 46.6 | 44.8 | 146.4* | 0.82 | 33 |
| Matched | 0.014 | 5.59 | 0.976 | 5.5 | 3.5 | 27.4* | 1.55 | 22 |

#### 3.4.5. Estimation of treatment effect on the treated

This part assesses the effect of the adoption of the Limu maize variety on the food security of rural households in order to meet the study’s stated objectives. The adoption of the Limu maize variety has had a significant and favorable influence on the food security of rural households, even after other features have been taken into account, according to the propensity score matching model with kernel matching. Adopter households of the Limu maize variety received 462.53 more calories per day per AE than non-adopter households (Table 8). Compared to non-adoptive households, adopter households consumed more calories on average. The fact that ATT is positive indicates that adopter households consume more calories than their counterparts. Accordingly, it was found that adopting the Limu maize variety had a positive effect on the food security of rural households. This finding agrees with (7, 33) which confirms positive relationship between technology adoption and farmers’ livelihood.

**Table 8:** Treatment effect on the treated

| Variable | Sample | Treated | Controls | Difference | S.E. | T-stat |
| --- | --- | --- | --- | --- | --- | --- |
| Daily calorie intake | Unmatched | 2555.868 | 2082.67 | 473.19 | 34.63 | 13.67 |
|  | ATT | 2566.91 | 2104.37 | 462.53 | 42.98 | 10.76 |

According to the study’s findings, households that adopted the Limu maize variety higher in terms of their food intake than non-adopter households. This results from the interactions between maize productivity, earnings, and food security. The maize variety Limu has been widely disseminated among farmers in the research area in an effort to increase maize yield. In this instance, more availability and access to food as well as marketable surplus for sale result from farmers’ higher maize yield. Additionally, household consumption can be indirectly increased by using the money made from the sale of surplus maize grain to buy other necessary food items. This suggests that the adoption of the Limu maize variety has a direct and favorable impact on household consumption, assuming other things remain constant. As a result, households in the study area that adopted the Limu maize variety have better food intake than households who did not.

One of the FGD participants claimed that he had experienced starvation five years prior. Even one piece of “injera” for his family was extremely difficult for him to obtain. He needs to borrow cash birr from his neighbors, popularly known as “*Arata*,” or he needs assistance from friends or family, particularly during the summer. Meanwhile, he was informed by one of his *Kebele’s* development agents that this major issue could be resolved by adopting more improved maize technology. After that, he made the decision to adopt improved maize seed to free himself and his family from this absurd situation. He then began utilizing Limu maize seed continuously after taking it. He could now produce 18 to 22 quintals of maize grain from a half hectare of land, making him a role model for farmers in his *Kebele*. He is gradually increasing his income to the point that he can now feed his family at least twice a day. His family ate corn as a “*injera*”. They occasionally provide porridge as a traditional cuisine as well as “kolo” and “Nefro” as fast food. In addition to its nutritional content, he used dried stack and cob as a source of energy for meal preparations as well as maize leaf and stalk for animal feed. They also purchase other foods by selling maize grain. This qualitative data from the FGD participants is consistent with the PSM model’s conclusions, which showed that adopters of the Limu maize variety had more food security than the variety’s non-adopters.

#### 3.4.6. Sensitivity analysis

PSM accounts for differences between treatment and control groups that can be seen, but it is vulnerable to differences that cannot be seen (34, 35). Different researchers become increasingly aware that is important to test the robustness of results to departures from the identifying assumption. Since it is not possible to estimate the magnitude of the selection bias with non-experimental data. This problem can be addressed by sensitivity analysis.

The final diagnostic step is a sensitivity analysis, which must be carried out to determine how sensitive the evaluated treatment impact is to unobserved factors that influence both treatment assignment and the outcome variable (30).

Rosenbaum (36) suggests utilizing the Rosenbaum bounding approach to assess the estimated ATT’s sensitivity with regard to deviation from the CIA. The impact of an unobserved variable is 0 if it has no impact on a particular study. As a result, only features that could be detected determined the participation probability. Even though two people have similar observed qualities, their chances of receiving the treatment may differ if there is unseen bias. This idea served as the foundation for the sensitivity analysis.

Table 9 demonstrates that, despite allowing adopter and non-adopter families to have different probabilities of being treated up to gamma=3.25(100%) in terms of unobserved factors, the adoption of the Limu maize variety had no impact on the food security of rural households. This suggests that even if we have set eγ mainly up to 3.25, which is a greater amount, the predicted ATT is still in doubt for outcome variables computed at different levels of critical value eγ. The impact estimated ATT of this study can therefore be inferred to be insensitive to hidden bias and to be a direct result of the adoption of the Limu maize variety.

**Table 9:**
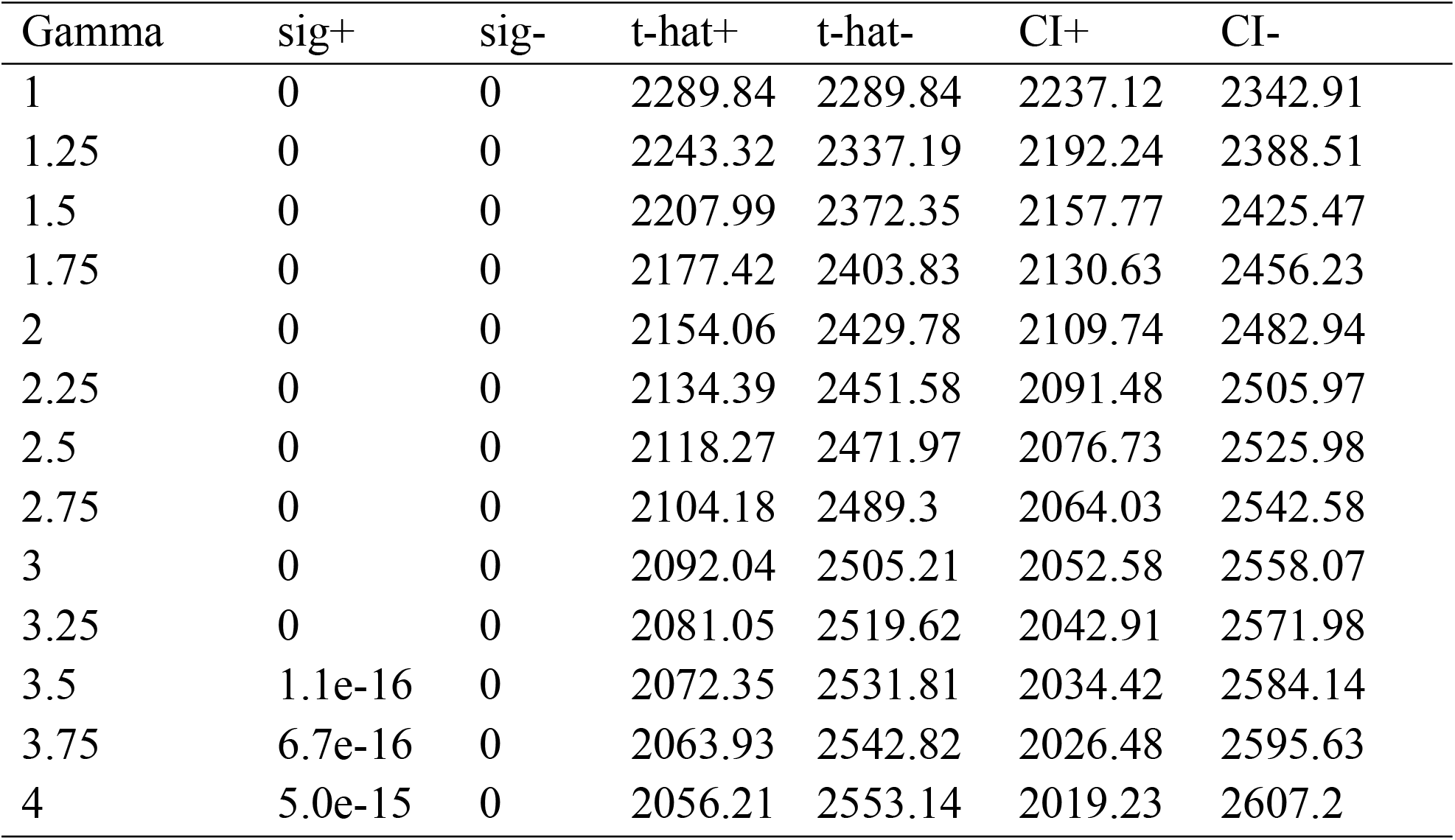
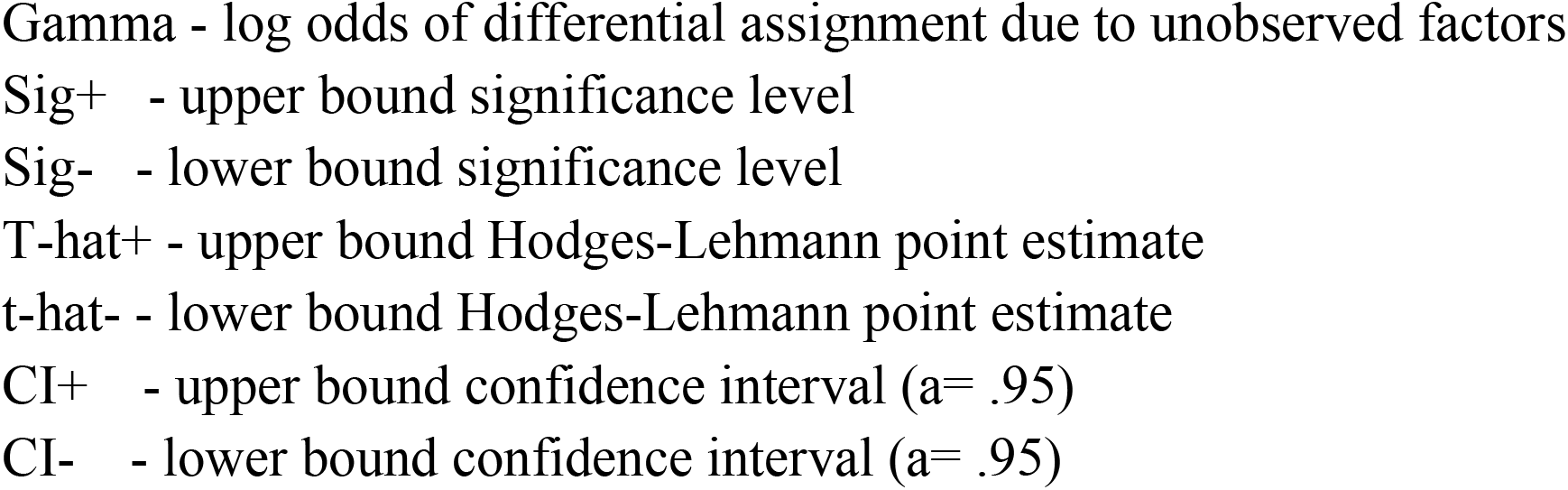
Result of sensitivity analysis using Rosenbaum bounding approach Rosenbaum bounds for KCal (N = 319 matched pairs) Rosenbaum bounds for KCal (N = 319 matched pairs)

## 4. Conclusion and recommendations

The determinants of adoption of the Limu maize variety and its effects on the food security of rural households in Ethiopia’s Dale Wabara district were the objectives of this study. To assess the collected data, both descriptive and econometric modeling techniques were used. The adoption of the Limu maize variety and its effects on the food security of rural households were both investigated using the econometric model’s binary logistic regression and PSM model, respectively.

Results from descriptive statistics showed that 48.9% of the sampled households adopted Limu maize variety, whereas 51.1% did not. Additionally, the results of a binary logistic regression model showed that the adoption of the Limu maize variety is positively influenced by factors like family size, household head’s education level, livestock ownership, total land size, the frequency of extension contact, and the availability of improved maize seed.

Additionally, using the propensity score matching model, the average treatment effect on treated is estimated based on the selection criteria for the matching method. The outcome supports the claim that using the Limu maize variety improves household food security. This indicates that, in terms of daily food intake at the household level, adopters of the Limu maize variety are much better off than non-adopters. In order to increase food security in the research area, it is advised that a greater distribution of the Limu maize variety be given priority. Additionally, it is important to enhance household adoption of the Limu maize variety in the research area in order to maintain the positive impact of Limu maize variety adoption. The authors also urge policymakers and planners to take into account the main elements that affect farmers’ decisions to adopt Limu maize varieties when developing their policies and plans for interventions aimed at improving agricultural production.

## Acknowledgements

At the outset, we would like to praise the everlasting Father and the Prince of love and peace the Almighty God who always let the bulk of unfinished work to be completed at a moment. Finally. We are grateful to all participants of this research.

## Availability of data

Data can be provided on request.

## Author contributions

Bikila, Adugna and Fikadu conceived the idea and designed the study; Bikila collected data and conducted the data analyses. Finally, Bikila, Adugna and Fikadu wrote the manuscript.

## Declarations

### Ethics approval and consent to participate

Approval to conduct the study was obtained from the Jimma University, college of agriculture and veterinary medicine. In addition, a permission letter was obtained from the Local Government Authority to collect data. In the villages, respondents were informed about the purpose of the study and consent to participate in the research was granted by each respondent before engaging them in the data collection process.

### Consent for publication

Final version of manuscript was approved for publication.

